# Inferring proteome dynamics during yeast cell cycle using gene expression data

**DOI:** 10.1101/170332

**Authors:** Krzysztof Kuchta, Joanna Towpik, Anna Biernacka, Jan Kutner, Andrzej Kudlicki, Krzysztof Ginalski, Maga Rowicka

## Abstract

Protein levels are most relevant physiologically, but measuring them genome-wide remains a challenge. In contrast, mRNA levels are much easier and less expensive to measure globally. Therefore, RNA levels are typically used to infer the corresponding protein levels. The steady-state condition (assumption that protein levels remain constant) is typically used to calculate protein abundances, as it is mathematically very convenient, even though it is often clear that it is not satisfied for proteins of interest. Here, we propose a simple, yet very effective, method to estimate genome wide protein abundances, which does not require the assumption that protein levels remain constant, and thus allows us to also predict proteome dynamics. Instead, we assume that the system returns to the baseline at the end of experiments; such an assumption is satisfied in many time-course experiments and in all periodic conditions (e.g. cell cycle). The approach only requires availability of gene expression and protein half-life data. As proof-of-concept, we calculated the predicted proteome dynamics for the budding yeast proteome during the cell cycle, which can be conveniently browsed online. The approach was validated experimentally by verifying that the predicted protein concentration changes were consistent with measurements for all proteins tested. Additionally, if proteomic data are also available, our approach can be used to predict how half-lives change in response to posttranslational regulation. We illustrated this application of our method with *de novo* prediction of changes in the degradation rate of Clb2 in response to post-translational modifications. The predicted changes were consistent with earlier observations in the literature.

## Introduction

Measuring protein abundance provides information which is not apparent from gene expression data and is crucial for the description of the state of a biological system (1). Nevertheless, measured mRNA concentrations are often used to linearly approximate the corresponding protein levels, even though it is known that such approximation will be very imprecise (1). Such indirect and inaccurate information is used because mRNA levels (unlike protein abundances) are relatively easy to determine due to RNA and DNA base pair complementarity, which enables precise and high-throughput measurements, such as sequencing and microarrays. Measuring protein levels remains challenging, due to the different chemical properties of proteins and wide dynamical range of protein abundances. Overall conclusion of the studies on the correlations between mRNA and the protein expression data (1-6) is that protein levels cannot be determined from mRNA levels just by correlation. Similar mRNA expression levels can be accompanied by a wide range (up to 20-fold difference) of protein abundances and vice versa (1). Relation between mRNA concentration, [*mRNA_i_*(*t*)], and protein concentration, [*P_i_*(*t*)], of the *i*-th protein can be described in the first approximation by a kinetic equation:

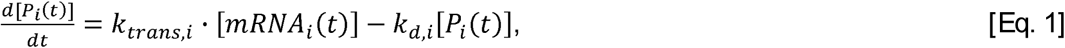

where 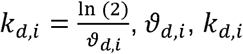 and *k*_*trans*,*i*_ are half-life, degradation rate and translation rate, respectively. Data regarding mRNA levels, protein abundances, degradation rates and translation rates are required to solve Eq. 1. Among these, only translation rates are not readily available for most model organisms. Eq. 1 is typically solved using the steady-state assumption, which is the easiest way to solve it mathematically, but it is also least physiologically relevant, since concentrations of many important proteins and their mRNAs are very dynamic. Therefore, we solve Eq. 1 in a new manner, instead of using steady-state assumption, we propose to use alternative boundary conditions. The alternative boundary condition we choose is that both mRNA and protein levels will be the same at time 0 and at the certain time T at the end of experiment. Such condition should be fulfilled in a typical control versus treatment experiment, at the time when treatment wears off as the cells go back to their original (control) state. Here, as proof-of-concept, we discuss a specific class of such experiments, where a system undergoes periodic changes, although periodicity of the data is not necessary to take advantage of our approach.

## Results and Discussions

Taking advantage of an availability of genome-wide data of mRNA levels, half-lives and average protein abundances in the model organism *S. cerevisiae*, we predicted dynamic protein abundances based on gene expression levels. We chose to use a simple, classical model of translation (7,8), which could be described by Eq. 1 above. The protein concentration [*P_i_* (*t*)] depends on the number of mRNAs ([*mRNA_i_*(*t*)]), which are translated with rate constant *k*_*trans*,*i*_, the protein-specific translation rate. Protein degradation is characterized by the rate constant 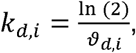 where *ϑ*_*d*, *i*_ is a protein half-life. The proposed model does not include variables reported sometimes to be proportional to the translation rates, such as ribosome occupancy or ribosome density (9). The reason, as we will show later, is that the minimal model, based only on data that is known with certainty to be relevant, performs better, as discussed below. Despite the simplicity of this model, it has been shown (10) that it may accurately capture the dynamical changes in protein abundances for a majority of human proteins. These results suggest that the model can be suitable for other eukaryotic systems (like *S. cerevisiae*) as well.

As described in detail in Materials and Methods, a protein concentration and its translation rate can be calculated from a time-course of its gene-expression measurements and its average abundance. As proof-of-concept, we chose five different *S. cerevisiae* cell cycle synchronized gene expression data sets (Table 1): alpha (3395 proteins), brd26 (2840 proteins), brd30 (2699 proteins), brd38 (2751 proteins), cdc15 (3173 proteins) and cdc28 (3424 proteins). First, we used periodogram to estimate consensus period for cell cycle periodically expressed genes in each of these data sets (Materials and Methods and Table 1). Second, we mathematically analyzed raw data on yeast protein half-lives, to remove negative values and improve overall accuracy of half-life estimates (Materials and Methods). Next, we used an existing compendium of the budding yeast mRNA and protein consensus levels to estimate these levels in our conditions (Materials and Methods). Finally, we numerically solved Eq. 1, using the Fixed Point Iteration method, for all periodically expressed proteins in these five data sets. This resulted in predicted time-courses of dynamical protein abundances, with 1-minute resolution during the whole cell cycle, for all budding yeast proteins available in each of five different data sets. All predicted dynamic protein concentrations and translations rates can be browsed, compared and downloaded via our web server (http://dynprot.cent.uw.edu.pl/).

**Table 1.**
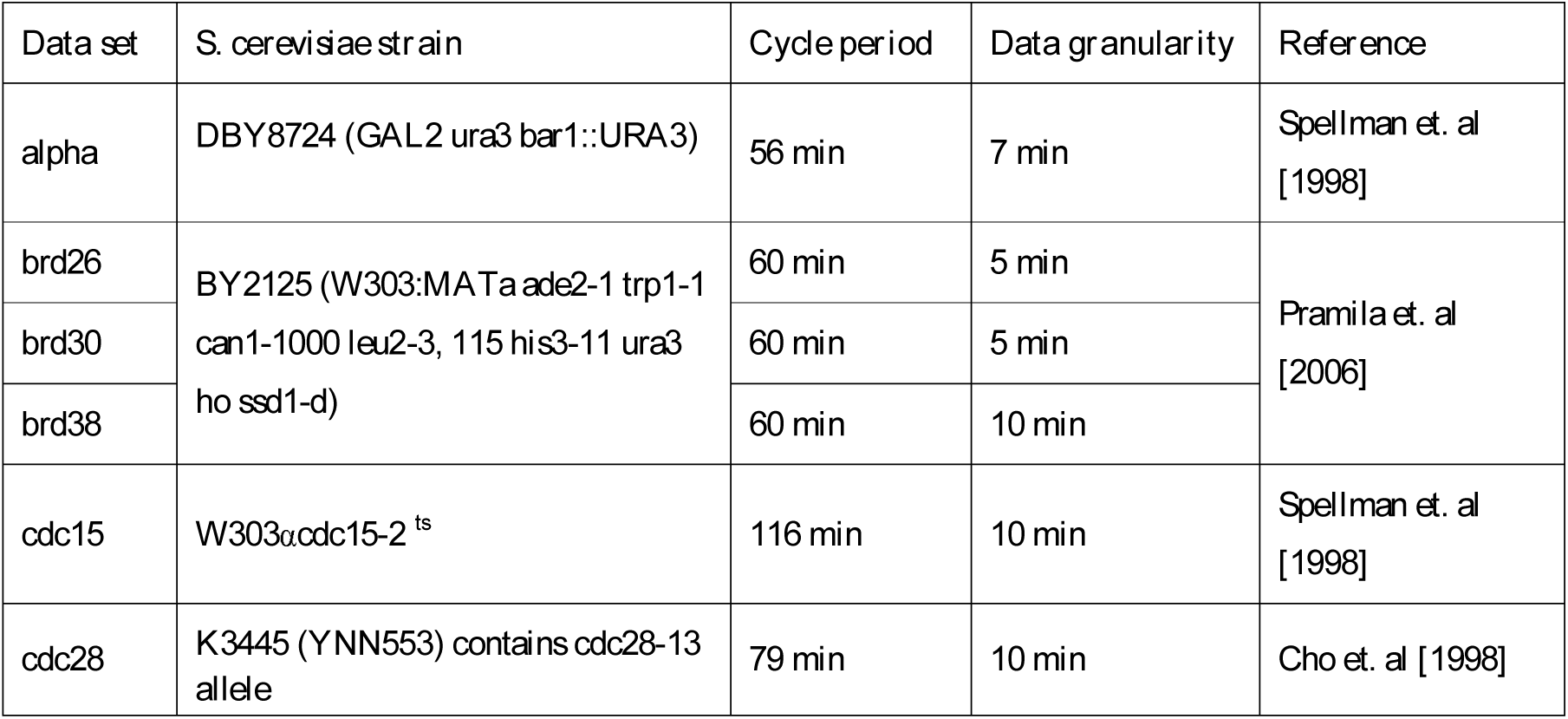
Data sources used.

### Validation of predicted dynamic protein abundances

In order to verify the temporal protein levels calculated using our model, we utilized western blotting to measure the actual protein concentrations for five representative proteins in cell cycle synchronized yeast culture (Materials and Methods). Representative proteins were chosen from the three groups: (1) proteins with relatively constant mRNA levels and predicted protein levels (Fig. 1A), (2) proteins with highly variable mRNA and relatively constant predicted protein levels (Fig. 1B), (3) proteins with variable mRNA and predicted protein levels during the cell cycle (Fig. 1C). For proteins with variable mRNA levels, we also required that they were transcriptionally regulated during the yeast cell cycle to guarantee that the observed changes in their levels would be meaningful; to confirm mRNA levels periodicity in the yeast cell cycle the SCEPTRANS web server was used (11). The choice of individual proteins within a group was based on availability of commercial antibodies. The first group is represented by Rad50p, the protein required for DNA damage repair, genetic recombination during meiosis and telomere maintenance (12,13). The levels of *RAD50* transcript remains almost constant during the cell cycle and due to a very long halflife of Rad50p (344 minutes, calculated as described in Materials and Methods using data of (14)) our model predicted that Rad50p levels should remain virtually constant during our experiments (Fig. 1A). Indeed, western blot analysis of the time-course Rad50p data confirmed this prediction (Fig. 2A). The second group is represented by histone Hht1 and Rnr1, the major isoform of the large subunit of ribonucleotide-diphosphate reductase, which is required for dNTPs synthesis (15). As these proteins are crucial for DNA replication, levels of their transcripts peak during S phase and decrease shortly afterwards. Despite this high variability of *HHT1* and *RNR1* transcripts, concentrations of their proteins during the cell cycle are predicted to be constant by our model due to the long half-lives of Hht1p and Rnr1p (349 and 77 min, respectively) (Materials and Methods, our analysis based on raw data of (14)). These predictions were confirmed by western blotting data showing no significant variability in the levels of Hht1and Rnr1 proteins during cell cycle progression (Fig. 2B). The last validation group consists of two proteins: Cdc5 and Clb2, which are directly involved in controlling cell cycle progression. Cdc5 is a polo-like kinase, necessary for meiotic progression (16), while Clb2 is a B-type cyclin required for transition from G2 to M phase (17). Their function is thus restricted to only specific stages of cell division. Consistent with this, both proteins are known to have transcripts levels strongly regulated during the cell cycle (11,18). According to our calculations based on O’Shea and colleagues data (14,19), Cdc5 and Clb2 half-lives are 10 and 22 min, respectively. Our model predicts that Cdc5 and Clb2 concentrations would exhibit strong variability during the yeast cell cycle (Fig. 1C). Indeed, the levels of Cdc5p and Clb2p as determined by western blotting vary strongly, reaching peaks at 65 and 115 min (M phase), and 55 and 110 min (G2/M transition), respectively (Fig. 2C). However, assuming that Clb2 has a constant half-life of 22 min (as calculated based on (14)), gives less than ideal agreement of predicted protein concentrations with western blot measurements (Fig. 2C).

**Fig. 1.**
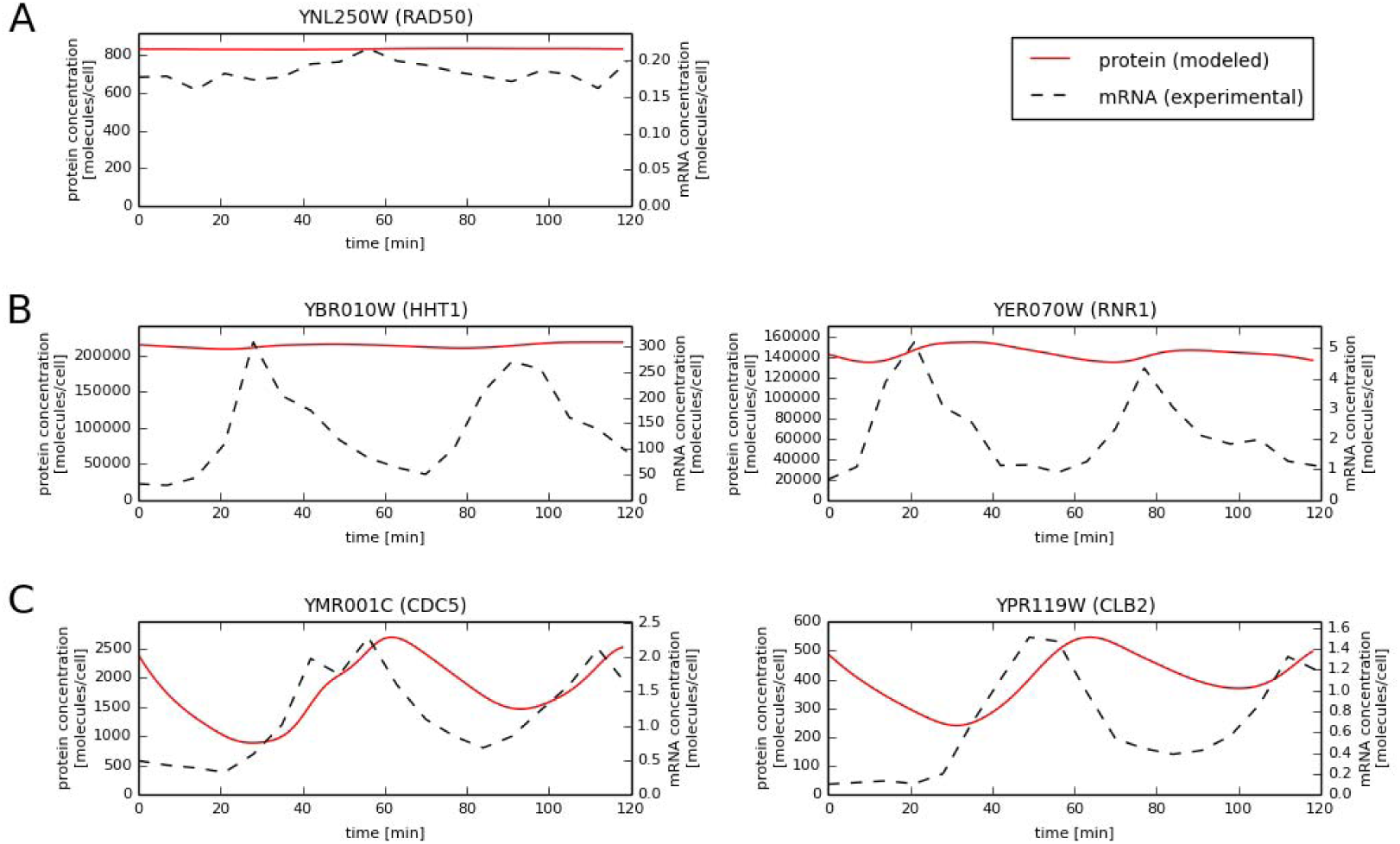
Comparison of mRNA vs. predicted protein concentrations for selected proteins in the alpha data set.

**Fig. 2.**
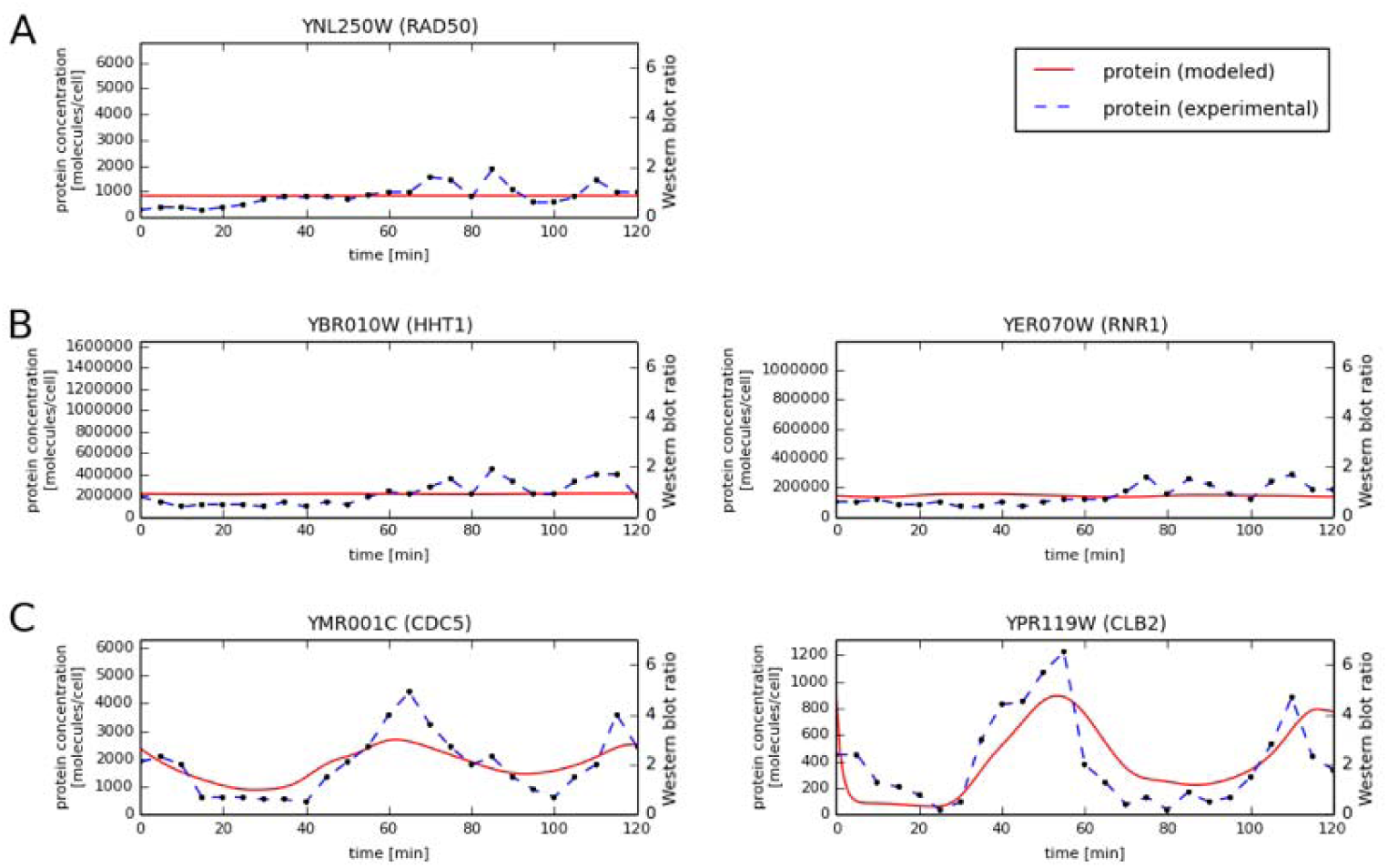
Comparison of experimental vs. predicted protein concentrations for selected proteins.

### Extension of the model to accommodate post-translational regulation

Discrepancies between predicted and experimental protein levels during the cell cycle may be caused by known inaccuracies of the western blot (up to 2-fold) or by post-translational regulation. To address this question, we also constructed a more complex model, allowing variable half-life throughout the cell cycle, to verify if considering dynamical half-lives would result in much better consistency of predictions with the experimental data. We tested the expanded model on the case of Clb2, since it was the only protein tested showing discrepancy with the predicted model beyond what would be expected from western blot measurement errors. The expanded model allows Clb2 to switch between longer and shorter half-lives depending on the stage of the cell cycle. We have generated such models for Clb2 with half-lives ranging from 1 to 40 minutes, with 1-minute step, and changing throughout the cell cycle. We chose the model which best fit the western blot data, which turned out to be the model assuming very short Clb2 half-life between the minutes 30 and 55 after the alpha-factor release and longer during the rest of the cell cycle (Fig. 3). Indeed, it was reported earlier that the Clb2 half-life was less than 1 min for cells arrested in G1 by α factor (14,19) and in our best-fitting models the Clb2 half-life was 1 minute (shorter values were not considered) during the G1 phase (Fig. 3). Clb2 had a longer half-life, closer to the value measured in (14,19) during the Clb2 activity window, which is at the G2/M transition. These results show another important application of our method: if half-life (and/or translation rates) are unavailable they can be estimated with good accuracy from corresponding gene expression and proteomic time-courses, even in very challenging cases in which half-life is variable and the protein time-course is inferred from relatively inaccurate western blots.

**Fig. 3.**
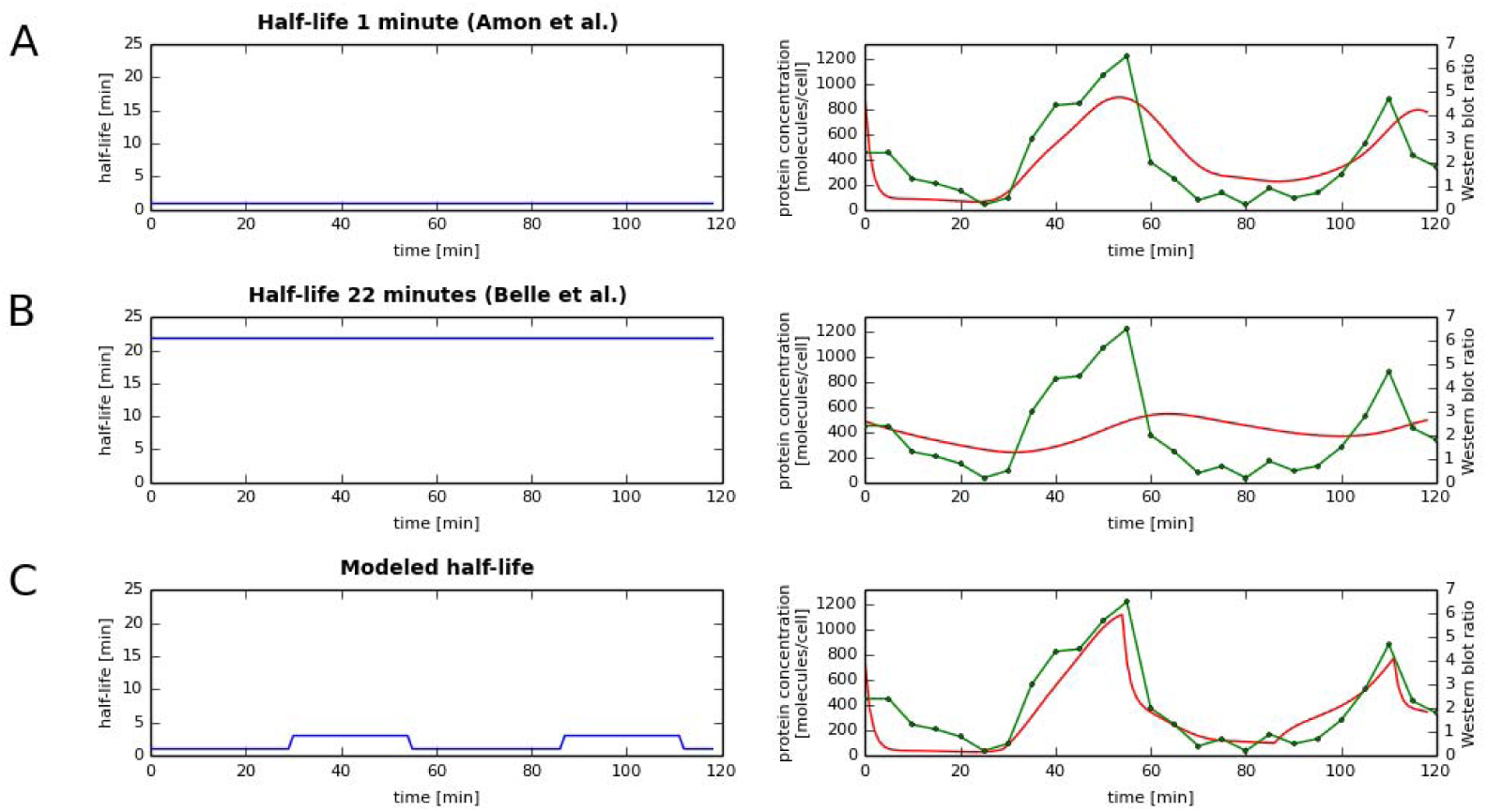
Variable half-life allows best fit of predicted (red) and experimentally measured (green) temporal protein concentration profiles. **A** and **B**: half-lives reported in the literature (left) for Clb2 does not lead to good fit of predicted and measured protein concentration temporal profiles (right), especially for half-life reported in (14) (**B**). **C**: variable half-life (left), found through numerical simulations (Material and Methods) allows for best fit between dynamic Clb2 abundances predicted from mRNA time-course and measured protein abundance time-course from the same condition.

### Correlation between mRNAs and protein abundances in time-course data

It is typically assumed that with an increase in quality of both gene-expression and proteomic data, the correlation between mRNA and protein abundance would grow. However, a significant correlation between mRNA and protein concentration can be expected only for some groups of proteins. Greenbaum et al. showed a significant increase in correlation between mRNA and proteins levels for proteins localized in the same cell compartment or with the same MIPS functional category (2). O’Shea and colleagues later showed that proteins of similar function tend to have similar half-lives. So far, the highest achieved correlations between mRNA and protein concentrations is the result of Futcher et al. (3), who found relatively high correlations (r=0.76) after transforming the data to normal distributions. The 0.7-0.8 range likely represents the highest correlation possible to achieve. On the other hand, protein half-lives are known to have a dynamic range of several orders of magnitude (14), and therefore even similar mRNA expression levels can be accompanied by a wide range of protein abundance levels, and vice versa (1). In general, it is increasingly recognized that mRNA abundances are only a weak surrogate for the corresponding protein concentrations mainly because of post-transcriptional control of gene expression. Our studies allow us to look deeper at this problem. We found that even though the Spearman and Pearson correlation between average protein and mRNA concentrations is highly significant (Table 2), temporal protein and mRNA concentrations are only weakly correlated (Fig. 4), with typical correlation not higher than 0.2. As expected, the highest correlations between temporary protein and mRNA abundances were observed for proteins with short half-lives, when proteins levels follow close behind mRNA concentrations (Fig. 5). These data show that even in the simplified case of not considering post-translational modification, mRNA levels are good estimates of temporal protein abundances during the whole cell cycle only for a handful of proteins, highlighting the usefulness of modeling them, as described above.

**Table 2.** Pearson and Spearman correlations between average mRNA and average protein concentrations.

| **Data set** | **Pearson** | **Person p-value** | **Spearman** | **Spearman p-value** |
| --- | --- | --- | --- | --- |
| alpha | 0.51 | 4.0e-224 | 0.58 | 2.3e-301 |
| brd26 | 0.43 | 2.1e-127 | 0.56 | 8.2e-236 |
| brd30 | 0.47 | 5.6e-148 | 0.57 | 3.0e-231 |
| brd38 | 0.53 | 4.0e-196 | 0.56 | 6.2e-227 |
| cdc15 | 0.52 | 1.1e-218 | 0.58 | 1.4e-279 |
| cdc28 | 0.52 | 2.0e-232 | 0.58 | 3.9e-302 |

**Fig. 4.**
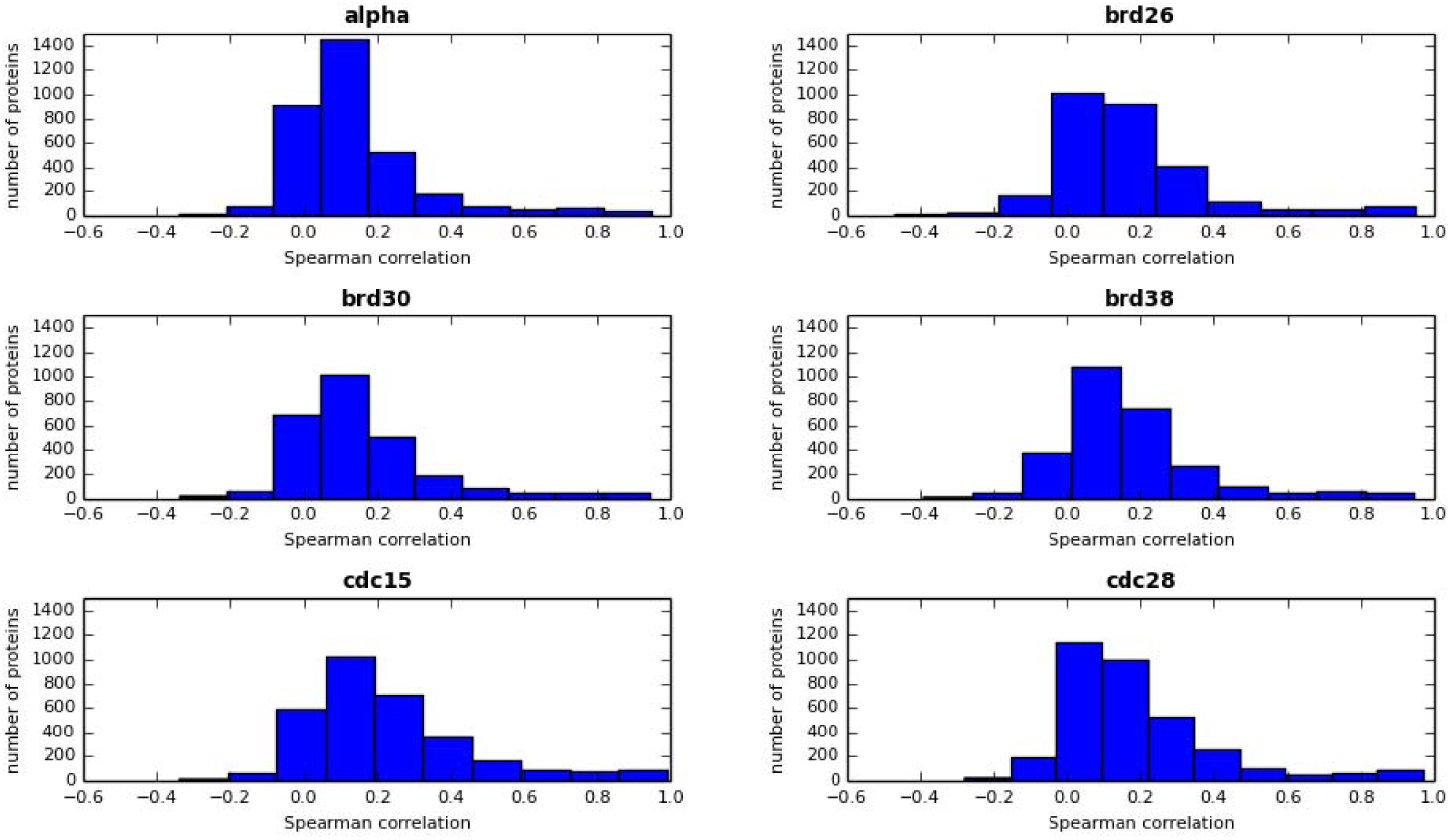
Histogram of the Spearman correlation between protein and mRNA concentrations during the cell cycle for all available proteins in the following data sets: alpha (3395 proteins), brd26 (2840 proteins), brd30 (2699 proteins), brd38 (2751 proteins), cdc15 (3173 proteins) and cdc28 (3424 proteins).

**Fig. 5.**
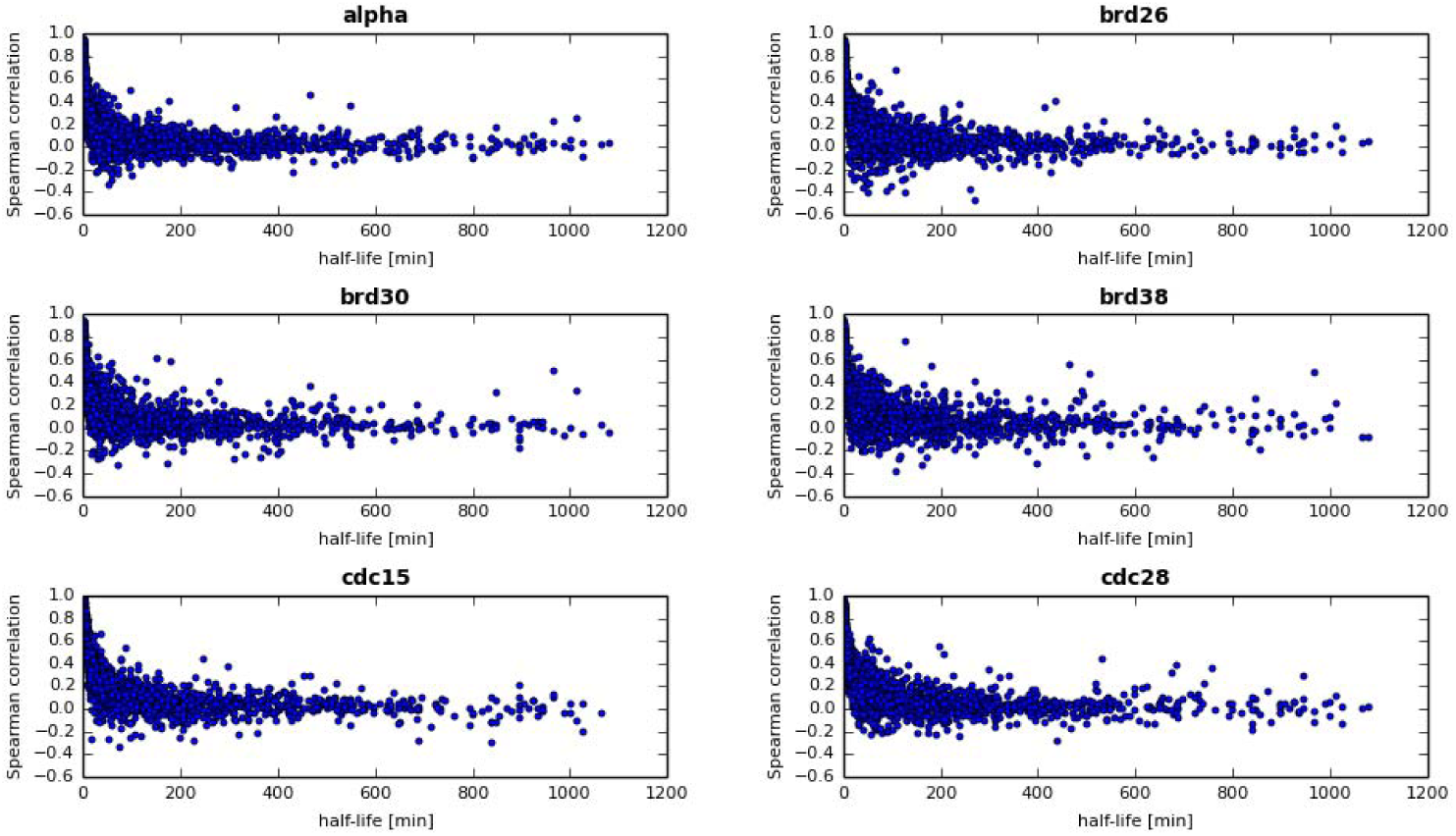
Relationship between Spearman correlations of protein and mRNA levels during the cell cycle and protein half-lives.

### Estimating translation rates

Translation rate (*TR*) is protein production rate (denoted by *k*_*trans*,*i*_ in Eq. 1). Translation rates are not easy to measure directly, and traditionally are estimated utilizing a steady-state condition (*TR_SS_*, Material and Methods, Eq. 7). However, steady-state condition is not typically fulfilled in physiological conditions. Moreover, there is a growing evidence that unlike degradation rate, translation rate is very plastic and is a mechanism to control protein abundances, in response to changing mRNA levels (e.g. (20)). Our approach provides a method for estimating condition-specific translation rate without requiring steady-state condition nor knowing protein abundance, but using time-series gene expression data instead (*TR_tc_*, Material and Methods, Eq. 2 and 3). To compare translation rates calculated using these two different approaches we computed *TR_diff_* (Materials and Methods, Eq. 6), which varies from 0 to 1 depending on how different *TR_ss_* and *TR_tc_* are. We found that there are relatively few proteins for which *TR_diff_* is greater than 0.1 (65 out of 3395 in the alpha data set, 59 out of 2840 in the brd26 data set, 25 out of 2699 in the brd30, 30 out of 2751 in brd38, 254 out of 3173 in cdc15 and 64 out of 3424 in cdc28). This result shows that our method offers a useful alternative to estimate translation rates when protein abundances are not known, but time-course genes expression data is available. We think the three main reasons for the observed discrepancies between these two methods of computing translation rates, which will be described in more detail below, are: (a) the effects of α-factor synchronization, (b) measurement errors of mRNA and protein concentrations and (c) time-dependence of halflives. (a) α-factor synchronization would cause mRNA levels of some genes to be changed, for example upon α-factor synchronization mRNA abundances of SST2/YLR452C (which regulates desensitization to α-factor (21)) and SW11/YGL028C (which may play a role in conjugation during mating based on its regulation by Ste12p (22)) are elevated. Indeed, for these two proteins we obtained *TR_diff_* > 0.1. (b) The second likely source of differences is measurement errors of data used: here mRNA and protein concentrations and degradation rates. (c) Third, as we will discuss below, some half-lives are time-dependent and neither steady-state nor the time-course based methods we used so far accommodate such time dependency. Due to very different manner of estimating *TR* using either the steady-state or time-course method it is not surprising that time-dependence of actual protein half-lives would affect these calculations in a different manner, causing observed discrepancies. In summary, main source of differences in translation rates we computed is related to our experimental conditions, with additional effects resulting from using time-course, not average expression values and from measurement errors.

*TR* was expected to correlate with many factors known to contribute to protein production, such as protein abundance, ribosome density, ribosome occupancy, mRNA concentration, codon adaptation index (CAI) or tRNA adaptation index (TAI) (23,24). However, the *TR_ss_* we computed (Fig. 6A) does not show high correlation with features expected to be correlated with it. For example, it seems intuitive and it has been proposed in Arava et al. (23) that *TR* would be proportional to ribosomal density (i.e. number of ribosomes bounded to mRNA) and ribosomal occupancy (number of mRNA associated with ribosomes) (their product is denoted by TA1 in Fig. 6A). However, there is no such correlation, neither Spearman nor Pearson (Fig. 6A). Although this could suggest that ribosomal density or occupancy do not contribute meaningfully to translation rates, the lack of high positive correlation between TR and proposed TR contributing factors is in fact the result of high standard deviations of 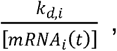 the proportionality factor between translation rate and average protein concentration for *i*-th protein. Indeed, the factors mentioned earlier, which are reported as likely to correlate with TR in some publications, are highly correlated with average protein concentration (Fig. 6B). TR is associated with average protein concentration, however, this correlation is not very high (0.18 for cdc15, 0.20 for brd30 and 0.19 for others) due to the important impact of half-life, which can vary by at least two orders of magnitude, on protein concentration (Eq. 8). Another interesting observation is that very complex attempts at modeling translation rate, such as Ribosomal Flow Model, do not fare better than simpler models: in our comparison complex RFM (24) performed worse.

**Fig. 6.**
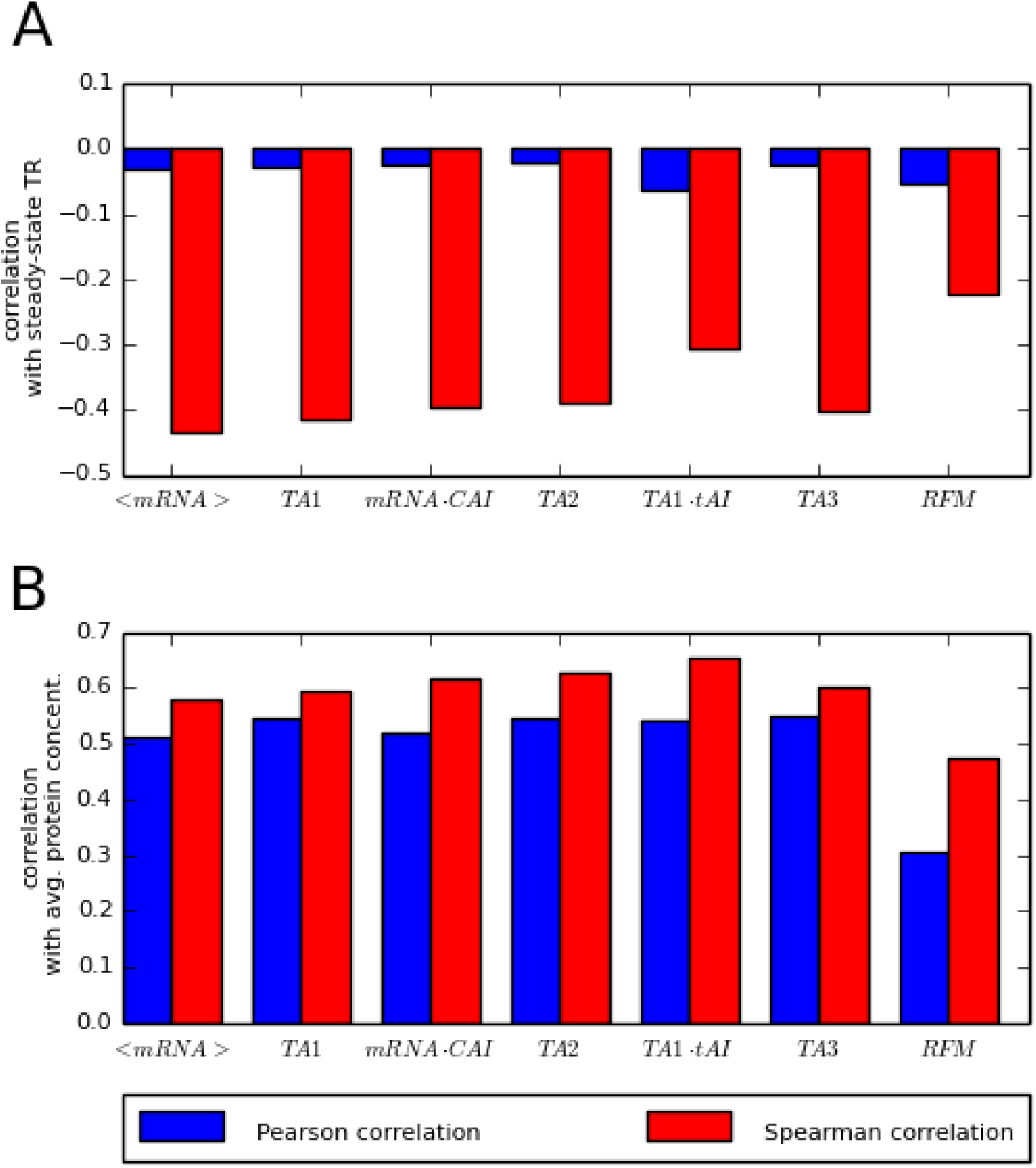
The Spearman (red bars) and Pearson (blue bars) correlations between: **A**: steady-state translation rates and translation rates descriptors, **B**: average proteins concentrations and translation rates descriptors. TA1, TA2 and TA3 were computed using the following formulae: TA1 = (ribosome density) ^∗^ (ribosome occupancy) ^∗^ (mRNA concentration), TA2 = (ribosome density) ^∗^ (ribosome occupancy) ^∗^ (mRNA concentration) ^∗^ CAI, TA3 = (ribosome density) ^∗^ (ribosome occupancy) ^∗^ (mRNA concentration) ^∗^ CAI/(0.06 + (ribosome density)) ^∗^ (ribosome occupancy ^∗^ mRNA concentration).

To visualize which cell compartments and protein functions are associated with high or low value of half-life and translation rate, we analyzed different MIPS functional categories and localizations using SCEPTRANS webserver (Fig. 7). Global analysis shows that half-lives and translation rates have almost the same levels in all functional categories. However, there are some interesting exceptions to this principle: in the cell wall and extracellular categories there are proteins with relatively short half-lives (that is high degradation rates) and high translation rates (Fig. B and D). Additionally, proteins involved in protein synthesis have much shorter half-lives than average (Fig. 7A).

**Fig. 7.**
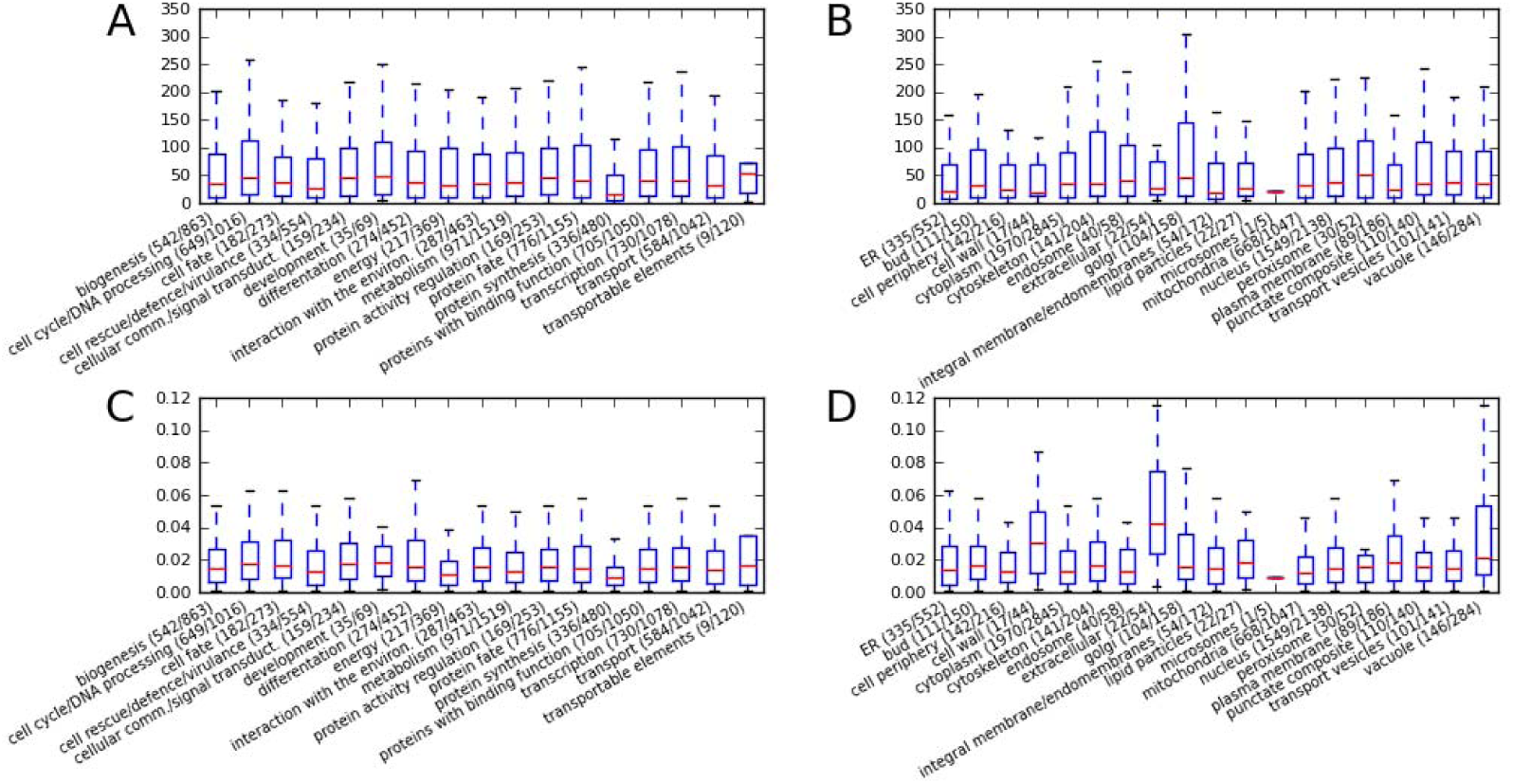
Half-lives (in minutes) (**A** and **B**) and translation rates (**C** and **D**) in each of the functional and localization categories as described in MIPS database, as retrieved from SCEPTRANS.

In summary, the proposed model (Eq. 1), combined with periodic data set (other time-course data sets can be used as well) allowed us to estimate not only genome wide changes in protein abundances, but also both translation and degradation rates of proteins. The model performs especially well in the most interesting case of substantial dynamic changes in protein abundances over time. It is also capable of detecting post-translational regulation of proteins for which corresponding time-course abundance data are available. Finally, calculated protein concentration time-courses were validated experimentally for several proteins.

## Conclusions

Taking advantage of the ready availability of genome-wide data of mRNA levels, we propose a model which predicts dynamic levels of protein abundances based on time-course of gene expression levels and measured or predicted half-lives. We experimentally verified the proposed computational approach in the model organism *S. cerevisiae* by measuring protein concentration changes for selected proteins in the α-factor synchronized cell cycle using western blotting. We also showed how our approach can be used to infer post-transcriptional or post-translational regulation, if both gene expression and proteomic time-course data are available. Additionally, we propose a method for estimating translation rates without using the standard, but typically non-physiological steady-state assumption. Instead, we propose to use a boundary condition of the beginning and end protein concentration equivalence, which is typically satisfied not only in periodic processes like the cell cycle, but also in common time-course experiments, when the system is allowed to return to baseline after treatment. Our approach may be useful in many experimental conditions where steady-state condition is clearly not satisfied, like in differentiation, but adaptive changes in translation rates play important regulatory role (20).

Motivation for our study was deeply practical: to provide estimated *in silico* time-course data for proteins for which corresponding gene expression measurements are known or to integrate genomic and proteomic data to elucidate possible post-translational regulation. Most other studies in the field were motivated instead by the desire to explain the observed degree of correlation between protein abundance and gene expression levels (1,2,25) or to estimate translation rates (24). Nevertheless, it seems that our estimation of translation rates – a necessary step on the path to estimate protein levels - is also rather accurate, perhaps more so than other popular methods (Fig. 6). Of course, in the case when proteomic data are unavailable, our predictions will be of limited accuracy for proteins undergoing posttranslational modifications and possibly additionally due to inaccuracies in the data measurement, especially half-lives (the half-life data we used in this study, (14), has multiplicative error of up to 2). Our goal, however, is not to produce accurate predictions for all proteins, but instead provide predictions that are far better than using mRNA as a proxy for a large number of proteins that are not highly unstable, but also do not undergo substantial post-translational regulation in the conditions studied. As was shown in our verification, and as should be expected, depending on half-life, protein abundance profiles may show anywhere between no resemblance, to very high resemblance to the underlying mRNA expression profiles. Therefore, our predicted protein profiles can be of valuable help for scientists interested in dynamic changes of protein abundances during their process of interest, but who only have gene expression profiles available, which are much easier and less expensive to measure than protein levels. Moreover, if a protein is known or predicted to undergo a post-translational modification, such as methylation (26) or phosphorylation (27), it can be flagged for potential lower accuracy of our predictions. If corresponding proteomic timecourse is available, potential temporal changes to half-life can be calculated, following the approach we used for Clb2. To allow such analysis in a variety of organisms and conditions, we are developing a webserver, based on the proof-of-concept study presented in this paper, to provide predicted protein time-course profiles based on user-provided gene expression and protein half-life data. Currently, all our predictions for proteome dynamics in the budding yeast in different conditions can be conveniently browsed and visualized at http://dynprot.cent.uw.edu.pl/.

In summary, we have shown that a simple model of the relationship between mRNA and protein levels usually leads to rather accurate prediction of protein levels, if post-translational regulation is not involved. Our approach can be used to obtain an approximate view of proteome dynamics (without post-translational regulation), to integrate gene expression and proteomic time-course data if both are available, or to more specific tasks, such as estimating changing degradation rates, as in our example with Clb2. Our approach was verified experimentally to provide useful results and we believe that such an approximated simulations of proteome dynamics may become a part of time-course gene expression analysis, either performed for the whole genome, or for pathways or genes of interest.

Recently, the availability of genome-wide measured protein degradation rates in various organisms (14,28) is growing (20,29), which makes our approach more broadly applicable. Moreover, there is also substantial progress in understanding how protein half-life is encoded in its sequence, which gives hope that these values may be predicted computationally from sequence alone in the coming years (30,31), which would allow the extension of our approach to any organism for which gene expression data are available.

## Materials and Methods

### Definitions

*Ribosome density* is an average number of ribosomes bound to mRNA per unit of mRNA length (100nt).

*Ribosome occupancy* is a fraction of transcripts associated with ribosomes, i.e. engaged in translation, with values in [0,1] interval.

### Quantitative model of gene expression

Using periodic gene expression data enables us to eliminate translation rate, *k*_*trans*,i,_ values from equation [Eq. 1]. In order to do that, we introduced function [*R*_*i*_(*t*)] defined as follow:

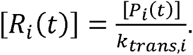

Since for a small change of time Δ*t*

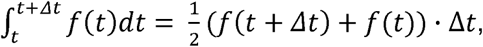

the first order differential equation [Eq. 1] can be rewritten in the form:

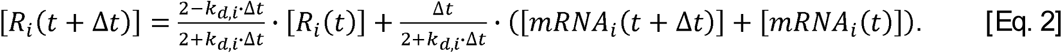

The boundary condition of the equation above:

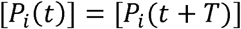

is equivalent to:

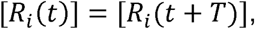

where T is the period of the cell cycle.

The proportionality factor *k*_*trans*,*i*_ can be obtained from a following formula:

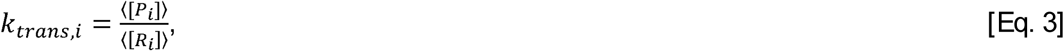

where 〈[*P_i_*]〉,〈[*R_i_*]〉 are the mean values [*P_i_*] and [*R_i_*] after time T, respectively.

### Data sets used

Average protein and mRNA concentrations have been taken from previous Beyer et al. studies (25). Test data sets alpha, brd26, brd30, brd38, cdc15 and cdc28 are cell-cycle synchronized gene expression data sets described in detail in Table 1. Data sets alpha and cdc15 have been published by Spellman et al. (32); cdc28 by Cho et al. (33) and brd26, brd30 and brd38 by Pramila et al. (34). The gene expression log_2_ ratios, *L_i_(t*), were transformed to mRNA concentrations [molecules/cell] in the following manner:

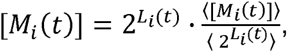

where 〈2^*L_i_*(*t*)^〉 is the arithmetic average of *2*^*L_i_*(*t*)^ in one cell cycle period and 〈[*M_i_*(*t*)]〉 is the cell-cycle average mRNA concentration in molecules per cell, based on literature (25). Linear interpolation was used to approximate a value of mRNA concentration in every minute during cell cycle, based on computed values at points of measurements (equation above).

### Estimating the consensus period for periodically expressed genes

The set of genes transcriptionally regulated during the cell cycle will be defined as the genes with a transcriptional modulation consistent with the periodicity *T* of the mitotic cell division. We utilized the measure of periodicity defined as the periodogram, *P*, (35-37) of transcript concentration:

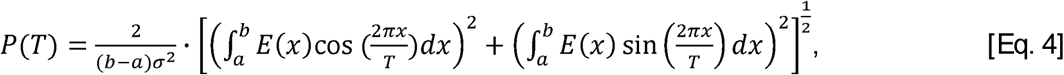

where *a* and *b* are the beginning and end of the time-course, respectively, function *E* is the transcript concentration and σ is the standard deviation of gene expression *E*. To accommodate uneven distribution of time points, we estimate *P*(*T*) using the unbiased formula of (36). The statistical significance of a single frequency (corresponding to periodicity with period T) in the periodogram, assuming a Gaussian null hypothesis, is expressed by

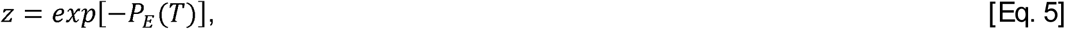

(35-38). Here, we do not have a reliable value of the period T measured independently from the transcriptome profiles. Therefore, similar as in (11,39), before applying Eq. 4, we estimated the most likely period of transcriptional oscillation in the system from the expression data. We have followed the Maximum Likelihood approach, using Eq. 4 and 5 for each gene independently over a range of possible periods, computing the logarithms of likelihood of periodicity for every gene and every period. These logarithms summed over all genes yield the total likelihood of every period, and the period with the maximum total likelihood has been adopted as the consensus period of regulation in the system. Estimated cell cycle periods for different data sets are described in Table 1.

### Correcting the estimated protein degradation half-lives

Belle et al. (14) reported protein half-lives, as estimated from the observed degradation rate, that sometimes have very high values, and, at times, negative ones. Since such values are not realistic, we adopted the following algorithm to estimate the most likely true half-lives for these proteins. We assumed that the measured quantity (degradation rate *k*_*d*,*i*_, which is related to half-life *ϑ*_*d*,*i*_ by 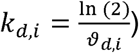 may include an error that has a Gaussian distribution, with a variance corresponding to the inverse of 300 minutes (the maximum reliably measureable value according to (14)) divided by the scaling factor *ln(2).* The negative reported half-lives result from experimental error, therefore, we used the described above error model and prior assumption that a half-life must be positive to correct the data. The true degradation rate was computed by integrating the normal distribution, limited and normalized to the positive part of its domain, and the inverse of this value multiplied by *ln(2*) was adopted as the corrected half-life. The correction was small for half-lives significantly shorter than 300 minutes, but significant for values longer than 300 minutes or negative reported values.

### Calculating protein concentrations

We used Fixed Point Iteration numerical method to solve Eq. 2 for each protein and mRNA data set. As a starting point for iterations we used [*R_i_* (0)] = 0 and Δ*t* = 1 minute. We continued iterative calculations until convergence, specifically until the condition |[*R_i_*(*T*)] – [*R_i_*(0)]| ≤ 5 · 10^−10^ had been met.

### Comparison between steady-state and time-course based translation rates

To determine the differences between steady-state derived translation rate, *TR_SS_*, and time-course derived translation rate, *TR_tc_*, we defined the coefficient *TR_diff_*:

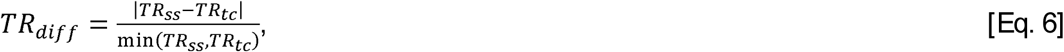

where time-course derived translation rate, *TR_tc_*, is defined by Eq. 3 and steady-state derived translation rate, *TR_ss_*, is defined as follows:

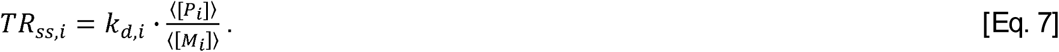

### Incorporating post-translational regulation

To accommodate post-translational regulation, we expanded our approach by allowing time-dependence of degradation rates. We will illustrate detecting post-translational modification discussing the example of Clb2. For Clb2, fitting constant degradation rate results in poor fit, both for half-lives based on O’Shea and colleagues report (14) (Fig. 3B) and for the much shorter half-life reported by Amon et al. (19) (Fig. 3A). Therefore, we propose instead a time-dependent half-life function that will be also periodic in the consecutive cell cycles. To describe a half-life that is modified by post-translational regulation within K minute window starting at the time *t_0_* within the cell cycle with the period T, we propose the following step function *ϑ*(*t*)*_d_*:

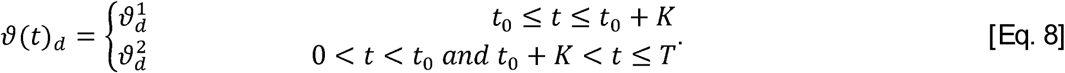

To find values of 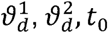 and K optimally describing time dependence of Clb2 half-life we numerically optimized these parameters, considering for half-lives 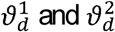 all values in the range from 1 minute to 40 minutes, with 1 minute step, and for *t*_0_ and K all possible times from the first to the last minute of the cell cycle, again with 1 minute step. For each set of parameters for the function *ϑ*(*t*)*_d_*, we solved the equation Eq. 2, as described previously (*Calculating protein concentrations*). The set of parameters offering the best fit with experimental data was chosen as the best estimate of true Clb2 half-life. Thus, we were also able to calculate the time-dependent degradation rate for Clb2 as 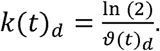 The best fit was achieved for variable half-life, with Clb2 protein becoming extremely unstable outside of the window of its activity during the cell cycle (Fig. 3C). This result shows that our approach allows one to re-discover, *ab initio*, the timing of post-translational regulation of a protein, if only gene expression and proteomic time-courses are available.

### α-Factor based synchronization

Yeast strain DBY8724 (Mat a *GAL2 ura3 bar1::URA3*) was kindly provided by P. T. Spellman. Obtained *S. cerevisiae* cells were synchronized by α-factor arrest as described by Spellman et al. (32) and later used by Pramila et al. (34). Cells were grown to an OD_600_ of 0.2 in YEP glucose pH 5.5, an asynchronous sample was taken and α-factor (Sigma Aldrich) was added to a concentration of 25 ng/ml. After 2 hours cells were released from α-factor arrest by pelleting and re-suspended in fresh medium to an OD_600_ of 0.2 (Fig. 8C, time 0). Every 5 min, for the next 120 min, 25 samples were taken (25 ml for western blot analysis, 1 ml for FACS analysis and 1 ml to count budding index). Cell cycle progression was monitored by bud counting and DNA content analysis (FACS) (Fig. 8A and 8B).

**Fig. 8.**
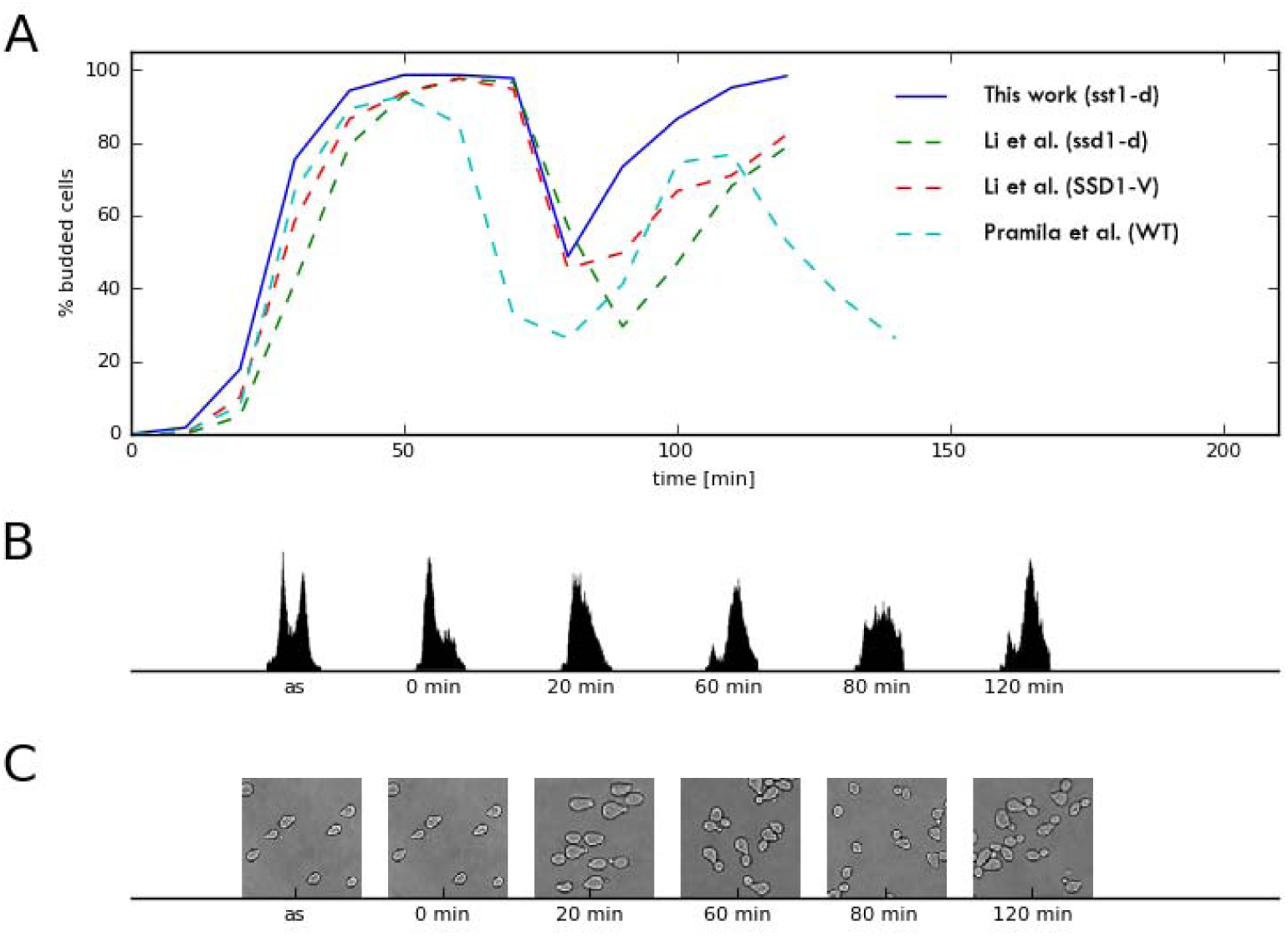
α-factor cell cycle synchronization. **A**: comparison of budding indices of our α-factor synchronization with those of Pramila et al. (34) and Li et al. (42), both for wild type (WT) and appropriate mutants. **B**: FACS results for asynchronous culture (as) and selected time points of our synchronization. **C**: yeast cells sampled from asynchronous culture and at selected time points.

### Budding index calculation and FACS analysis

For budding index calculation, two hundred cells were examined at every time point. The budding percentage was calculated as the number of budded cells divided by the number of all cells. To monitor the DNA synthesis, samples were prepared as described previously (40) and DNA content was measured using a BD FACSCalibur Flow Cytometer.

### Western blot analysis

Cell extracts were prepared by TCA precipitation (41) and then subjected to the western blot analysis. Protein samples were separated on Mini-PROTEAN TGX 4–20% (Bio-Rad) gels and transferred to PureNitrocellulose Paper 0.45 μm (Bio-Rad). Blots were blocked using 0.2% I-Block buffer (Applied Biosystems), cut horizontally and probed with primary antibodies followed by incubation with appropriate horseradish peroxidase-conjugated secondary antibodies. The primary antisera used to detect selected proteins were from Santa Cruz Biotechnology (Rad50, Cdc5, and Clb2), Abcam (H3), Agrisera (Rnr1) and Millipore (Act1) and the secondary antisera were from Dako. Protein bands were visualized with the Immoblilon Western (Millipore) and scanned in G-Box imaging system (Syngene). Band intensities were quantified using Gene-Snap software (Syngene).

## Acknowledgments

This work was supported by Foundation for Polish Science (TEAM), National Science Centre (2011/02/A/NZ2/00014, 2014/15/B/NZ1/03357), European Regional Development Fund under Innovative Economy Programme (POIG.02.02.00-14-024/08-00, POIG.02.03.00-14-128/13) and NIH NHLBI grant NHHSN268201000037C (N01-HV-00245) to UTMB-NHLBI Proteomic Center for Airway Inflammation and by NIH NIGMS grant R01 GM112131. We are grateful to Andrzej Dziembowski and Maciej Sykulski for insightful suggestions, Paul T. Spellman for DBY8724 *S. cerevisiae* strain and help with α-factor based synchronization, and Tamir Tuller for RFM translation rates data.

